# An optimized procedure for robust volitional drug intake in mice

**DOI:** 10.1101/786616

**Authors:** Alberto J. López, Amy R. Johnson, Ansley J. Kunnath, Jennifer E. Zachry, Kimberly C. Thibeault, Munir G. Kutlu, Cody A. Siciliano, Erin S. Calipari

## Abstract

Substance use disorder is a behavioral disorder characterized by volitional drug consumption, compulsive behavior, drug seeking, and relapse. Mouse models of substance use disorder allow for the use of molecular, genetic, and circuit level tools, which provide enormous potential for defining the underlying mechanisms of this disorder. However, the relevance of results depends entirely on the validity of the mouse models used. Self-administration models have long been considered the gold standard of preclinical addiction models, as they allow for volitional drug use, this providing strong face validity. In a series of experiments, we show that traditional mouse models of self-administration, where behavior is maintained on a fixed-ratio one schedule of reinforcement, show similar levels of responding in the presence and absence of drug delivery - demonstrating that it is impossible to determine when intake is and is not volitional. Further, when assessing inclusion criteria, we find a sex-bias in exclusion criteria where females that acquired food self-administration were eliminated when traditional criteria were applied. To address these issues, we have developed a novel mouse self-administration procedure where animals do not need to be pre-trained on food and behavior is maintained on a variable ratio schedule of reinforcement. This procedure increases rates of reinforcement behavior, increases levels of drug intake, and eliminates sex bias in inclusion criteria. Together, these data highlight a major issue with fixed-ratio models in mice that complicates subsequent analysis and provide a simple and novel approach to minimize these confounds with escalating variable-ratio schedules of reinforcement.

## Introduction

A sizable effort has focused on creating mouse models of substance use disorders (SUD) to define the molecular and circuit-based mechanisms that underlie addictive behaviors. Intravenous self-administration is the “gold standard” for preclinical models of SUD because it allows for the complex assessment of multiple components of volitional drug intake and provides an animal model for changes observed in humans throughout the transition to SUD[1–10]. The development of mouse self-administration models along with novel molecular, circuit, and genetic tools has allowed for the expansion of our understanding of the underpinnings of these behaviors with unprecedented resolution[11–16]. However, in many cases behaviors that have been validated in other model systems have been applied to mice without considering the different factors that influence these behavioral read-outs across species[17]. Here, we conducted a series of studies to empirically evaluate interpretations from existing mouse self-administration models which are standard in the field. We then used these evaluations to establish a novel mouse self-administration procedure optimized to increase drug intake, produce more consistent responding, and eliminate biases in inclusion criteria.

Volitional drug consumption is a core feature that makes self-administration models more translationally relevant than experimenter-delivered injections[4,18–25]. However, generating robust operant self-administration models in mice has been challenging. Currently, mouse models commonly rely on food pre-training where responses on the active lever results in the delivery of food with a conditioned cue[5–7,25]. In subsequent sessions the reinforcer is switched to drug, representing a contingency switch, where the mouse now learns that the same response results in a different outcome. Typically, if responding continues on the active lever after the contingency is switched to drug, mice are said to have learned/acquired the self-administration task and are included in the study. Confounding this interpretation is the fact that the previous food training may produce cue-food association, and these conditioned cues may be capable of supporting robust responding on their own[26–28]. Further, neutral cues alone can support reinforcement, even on high effort schedules in mice[29–31]. Thus, responding on the active operanda could be maintained by the cues, conditioned reinforcement, or lack of food response extinction – rather than being maintained by the delivery of drug. These factors make it difficult to ascertain if self-administration behavior is truly volitional, and without clear controls to interpret these behaviors, setting empirically derived acquisition/inclusion criteria is impossible.

Here we sought to identify the core factors that underlie operant behavior in mice and to develop a mouse training paradigm that does not require food training, reduces biases in inclusion criteria, and results in higher intake with stable rates of behavior. First, we find that traditional training criteria are particularly problematic in mice where they often meet these criteria in the absence of a reinforcer. Further, inclusion rates for male and female subjects were differentially impacted by using standard acquisition criteria to interpret these behaviors. We then developed a simple procedure that relies on variable-ratio schedules of reinforcement (VR) which eliminates the need for food pre-training as well as sex-bias in inclusion criteria. Moving forward, this standardized paradigm will be effective in further characterizing the neural mechanisms underlying drug-seeking behavior.

## Methods

### Animals

8-week old male and female C57BL/6J mice were purchased from the Jackson Laboratory and maintained in a 12-hour 6:00/6:00 reverse dark/light cycle with food and water provided ad libitum. During self-administration, animals were food restricted to ∼95% body weight with water provided ad libitum. Experiments were approved by the Institutional Animal Care and Use Committee of Vanderbilt University Medical Center. All experiments were conducted according to the National Institutes of Health guidelines for animal care and use.

### Jugular catheter implantation

Mice were anesthetized with ketamine (100 mg/kg) and xylazine (10 mg/kg) IP and implanted with chronic indwelling jugular catheters, as previously described[10,32]. Catheters were custom made and consisted of a back-mounted pedestal with silicone tubing (Access Technologies #BC-2S; 0.3mmID, 0.6mmOD) and a silicone bead was placed 1 cm from the end of the tubing as an anchor to suture the catheter into the vein once implanted. Ampicillin (0.5mg/kg)/heparin (10U/mL) in 0.9% saline was administered IV daily. Mice recovered >3 d before commencing training.

#### Self-Administration

Mice were trained in standard operant chambers (Med Associates, St Albans, USA) equipped with 2 illuminated nose-pokes, and a white noise generator with speaker. During each daily session, the initiation of white noise signaled the beginning of, and remained on throughout, the session. For each task, one nose-poke was designated as the “active poke” that would result in the delivery of the reinforcer and the other nose-poke was the “inactive poke” and had no delivered reinforcer, but program consequences depended on each experiment as outlined below.

### Sucrose self-administration and conditioned reinforcement (Figures 1 and 2)

Active nose-pokes resulted in the delivery of a single, 45 mg, sucrose pellet (Dustless Precision Pellets, Bio-Serv F0025, chocolate flavor) paired with a nose-poke light illumination (5 sec), on a fixed-ratio 1 (FR1) schedule. Inactive nose-pokes had no programmed consequences. Following 3 consecutive days of FR1 training, mice were trained for 3 days under an FR2 schedule as described previously[5]. We switched animals from food self-administration to conditioned reinforcement where mice responded for just the delivery of the 5 sec nose-poke light, in the absence of any reinforcer.

**Figure 1:**
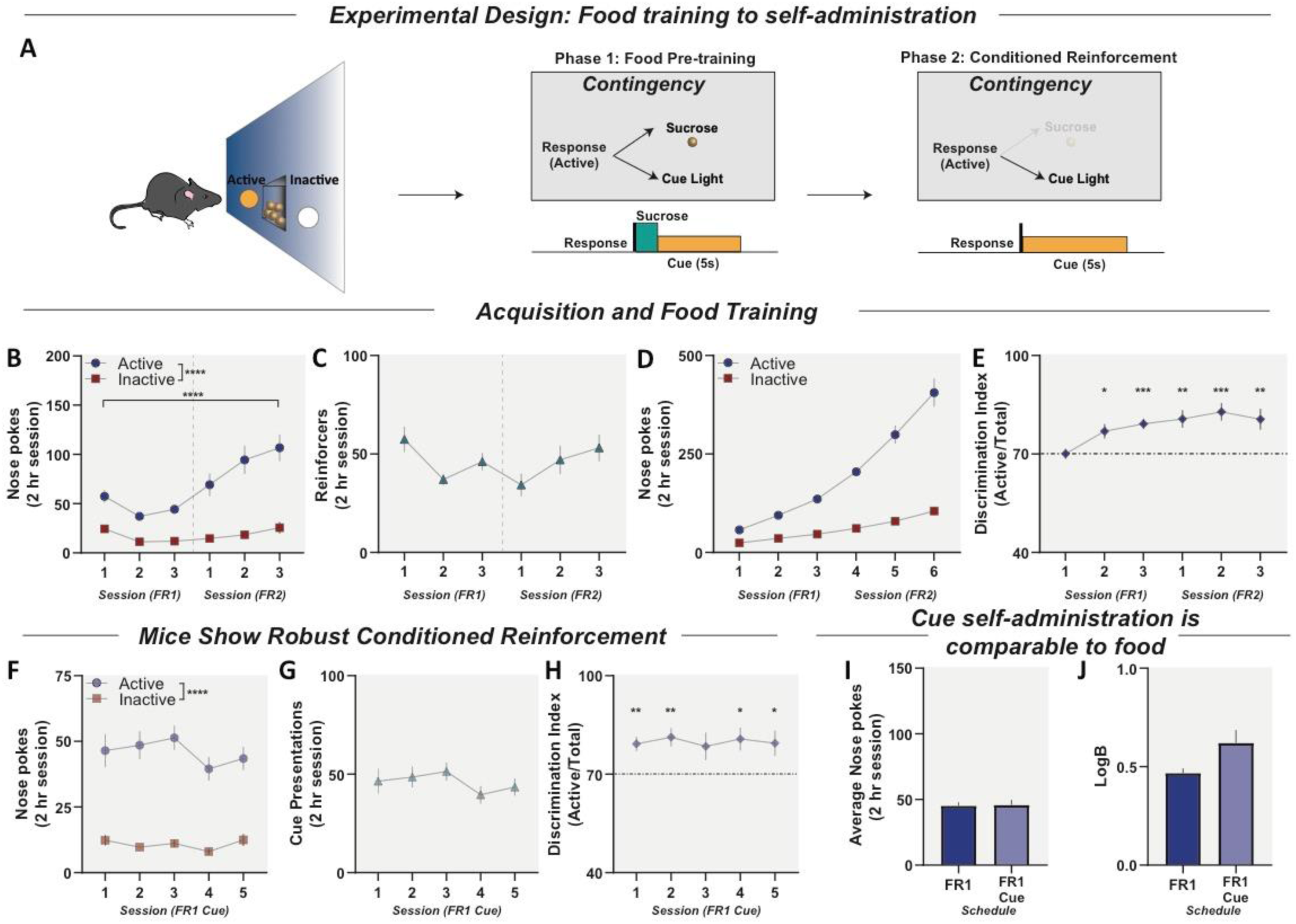
Typical pre-training approaches confound drug self-administration experiments in mice. Previous food-paired cues support conditioned reinforcement. (**A**) Illustration of operant training procedures. Phase 1, Operant Training: Active nose-pokes delivered a sucrose pellet concurrent with a 5 sec cue light on a fixed ratio one (FR1) and a fixed ratio two (FR2) schedule of reinforcement. Phase 2: Conditioned Reinforcement. Mice were switched from food responding to an FR1 schedule -like would be seen in cocaine self-administration training -however, active pokes resulted in only a 5 sec cue light presentation, with no reinforcer. In all phases, inactive nose-pokes were recorded, but had no programmed consequence. (**B**) Active and inactive nose-pokes during FR1 and FR2 food training. Mice show higher responding on the active lever compared to inactive and increase active responding during FR2. (**C**) Sucrose pellets earned across training. Animals earn reinforcers at a stable rate across FR1 and FR2 schedulesof reinforcement. (**D**) Cumulative record of active and inactive responses across training sessions. (**E**) Discrimination index across food training plotted as active/total responses. (**F**) Active and inactive nose-pokes during FR1 during conditioned reinforcement. Mice maintain active responding for the presentation of the cue alone for 5 consecutive days. (**G**) Cue presentations earned across sessions. Animals earn stable cue presentations over 5 consecutive days. (**H**) Discrimination index across cue testing showing that mice still meet discrimination criteria with no reinforcer present. (**I**) Number of active nose-pokes for food or for cues alone (with no reinforcer present) are indistinguishable under an FR1 schedule. (**J**) Mice show similar bias (measured by Log *b*)for active lever when reinforced by food or cue light presentations alone. Data reported as mean ± S.E.M. * p < 0.05, ** p < 0.01, *** p < 0.001, **** p < 0.0001

**Figure 2:**
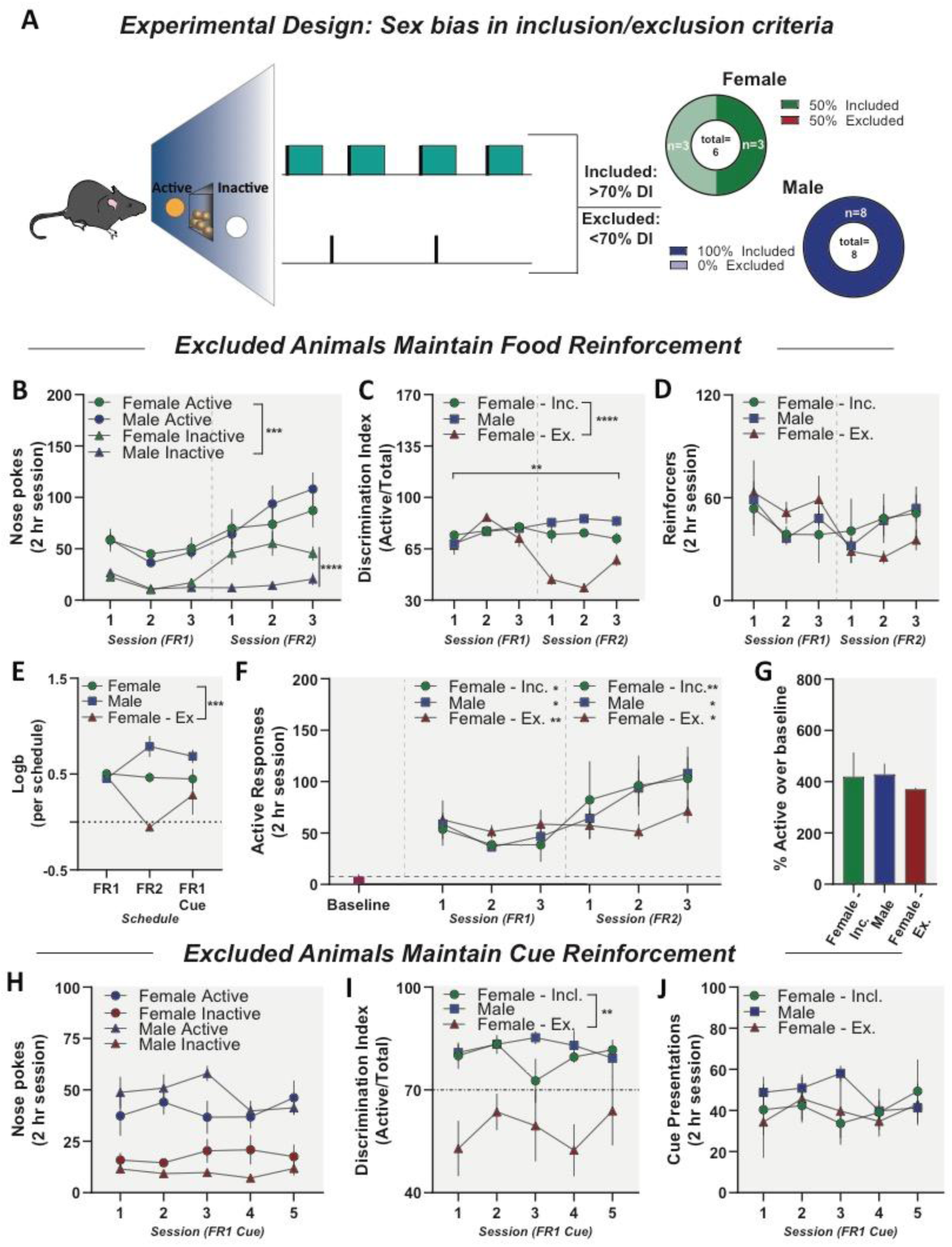
Conventional operant criteria create a sex bias. (**A**, *left*) Illustration of traditional food pre-training criteria, where animals must meet a discrimination index 70% active responses. (**A**, *right*) Criteria excludes more female than male mice (**B**) Active and inactive responses for males and females over training sessions. Females increase inactive responding under FR2, while both males and females increase active responding under FR2 (**C**) Discrimination indices of Male, Included Female, and Excluded Female mice over sucrose training sessions. Excluded Females show decreased discrimination index over time compared to Included Female and Male mice. (**D**) Food pellets earned across food training. Male, Included Female, and Excluded Female mice earn a similar number of food reinforcers. (**E**) Included animals maintain bias for active responding across training and test schedules, while Excluded Females show a decrease in bias for active nose-poke. (**F**) Because inactive nose-poke responses have no programmed consequences, another way of assessing reinforcement is a change in behavior from baseline rates of responding. Untrained mice show low baseline levels of responding on the “Active” nose-poke. All mice respond above unreinforced baseline response levels under (**F**, *middle*) FR1 and (**F**, *right*) FR2 schedules of food reinforcement. (**G**)All groups show comparable levels of responding for sucrose above baseline. (**H**) Active and inactive responses for males and females over cue testing sessions. (**I**) Discrimination indices of Male, Included Female, and Excluded Female mice over cue testing sessions. Excluded Females show decreased discrimination index during cue testing (**J**) Male, Included Female, and Excluded Female mice maintain similar levels of cue presentations during cue testing indicating that female removal is due to high inactive responses, not a lack of acqusition. Data reported as mean ± S.E.M. * p < 0.05, ** p < 0.01, *** p < 0.001, **** p < 0.0001

#### Baseline Responding

Mice were placed into the chamber and nose-pokes on either operanda were recorded but had no programmed consequence over three consecutive days. The average number of responses per session is reported as the baseline rate of nose-poking behavior.

### Variable ratio self-administration paradigm in mice (Figures 3 and 4)

Animals underwent intravenous cocaine self-administration where active nose-pokes resulted in a single infusion of cocaine (1 mg/kg/infusion; 0.035ml, 3 sec) or sterile saline (0.9% NaCl; 0.035ml, 3 sec) and a concurrent 5 sec delivery of active nose-poke light. Reinforced responses on the active nose-poke initiated a 5 second timeout (concurrent with reinforcer + cue delivery). Inactive nose-pokes resulted in inactive nose-poke light delivery with the same parameters as the active lever.

**Figure 3:**
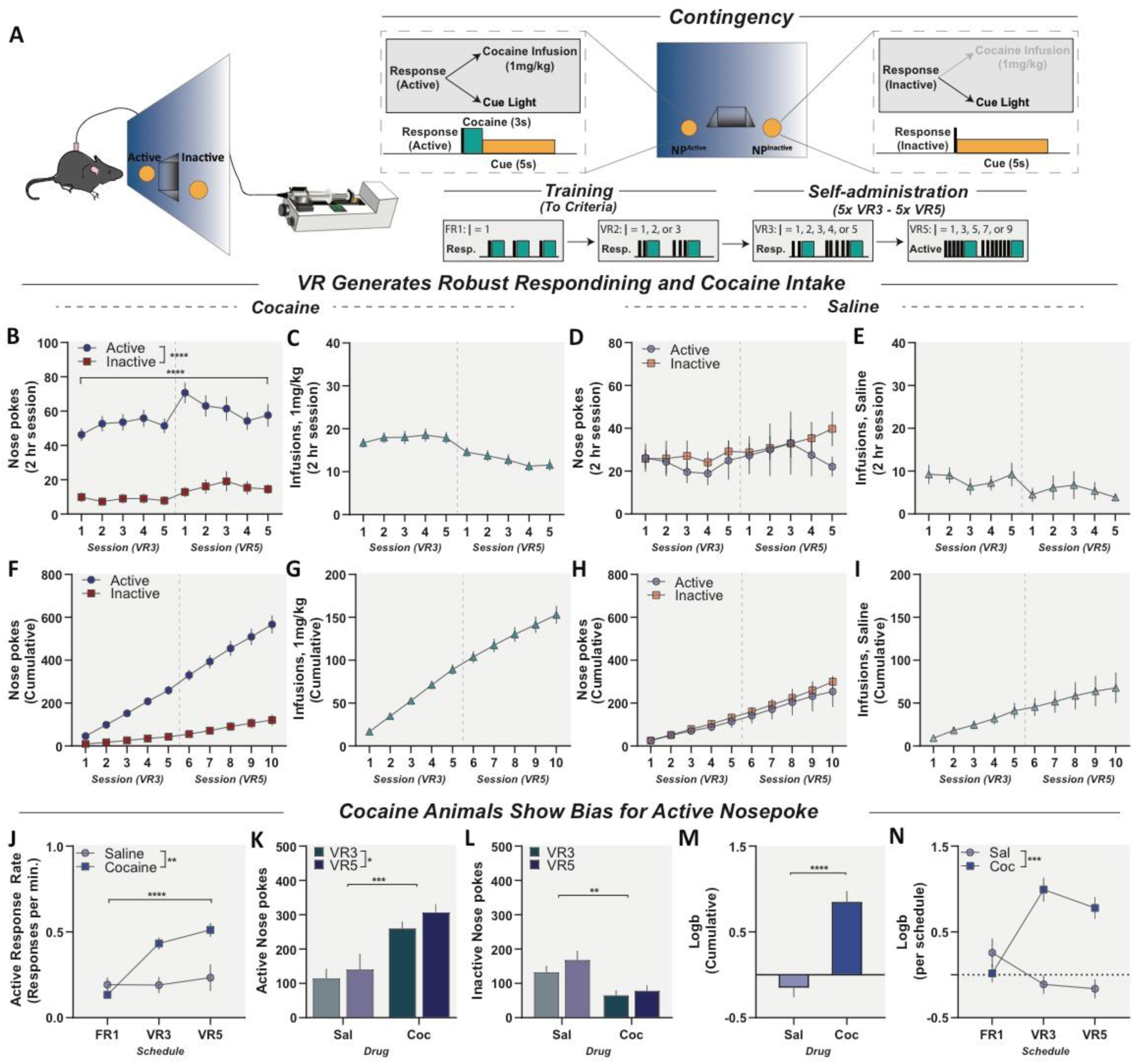
New procedure for drug self-administration in mice that requires no food pre-training, shows robust self-administration, and increases drug intake. (**A**, *left*) Schematic for mouse intravenous cocaine self-administration. (**A**, *right*) Training and Self-administration schedule. (**B**) Active and inactive responses for cocaine (1mg/kg/inj) self-administration in mice on VR3 and VR5 schedules of reinforcement. Animals show a preference for active nose-poke compared to inactive that is schedule dependent. (**C**) Total cocaine infusions per session. (**D**) Active and inactive responses for saline in mice on VR3 and VR5 schedules of reinforcement. Mice do not show a preference for either the active or inactive lever. (**E**) Total saline infusions per session. (**F**) Cumulative record of active and inactive responses across cocaine self-administration. (**G**) Cumulative record of cocaine infusions acrosscocaine self-administration. (**H**) Cumulative record of active and inactive responses across saline self-administration. (**I**) Cumulative record of saline infusions across saline self-administration. (**J**) Response rates on the active poke throughout training and self-administration. Cocaine animals demonstrate increased response rates on the active lever throughout cocaine self-administration compared to saline controls. (**K**) Total active responses for saline and cocaine animals during VR3 and VR5. Cocaine animals respond more on active poke and increase responding under VR5 compared to saline controls. (**L**) Total inactive responses for saline and cocaine animals during VR3 and VR5. Cocaine animals respond less on the inactive poke compared to saline controls. (**M**)Cocaine animals demonstrate increased bias for active nose-poke during VR self-administration compared to saline controls. (**N**) Cocaine animals show increased bias during VR self-administration compared to saline controls. Data reported as mean ± S.E.M. * p < 0.05, ** p < 0.01, *** p < 0.001, **** p < 0.0001

**Figure 4:**
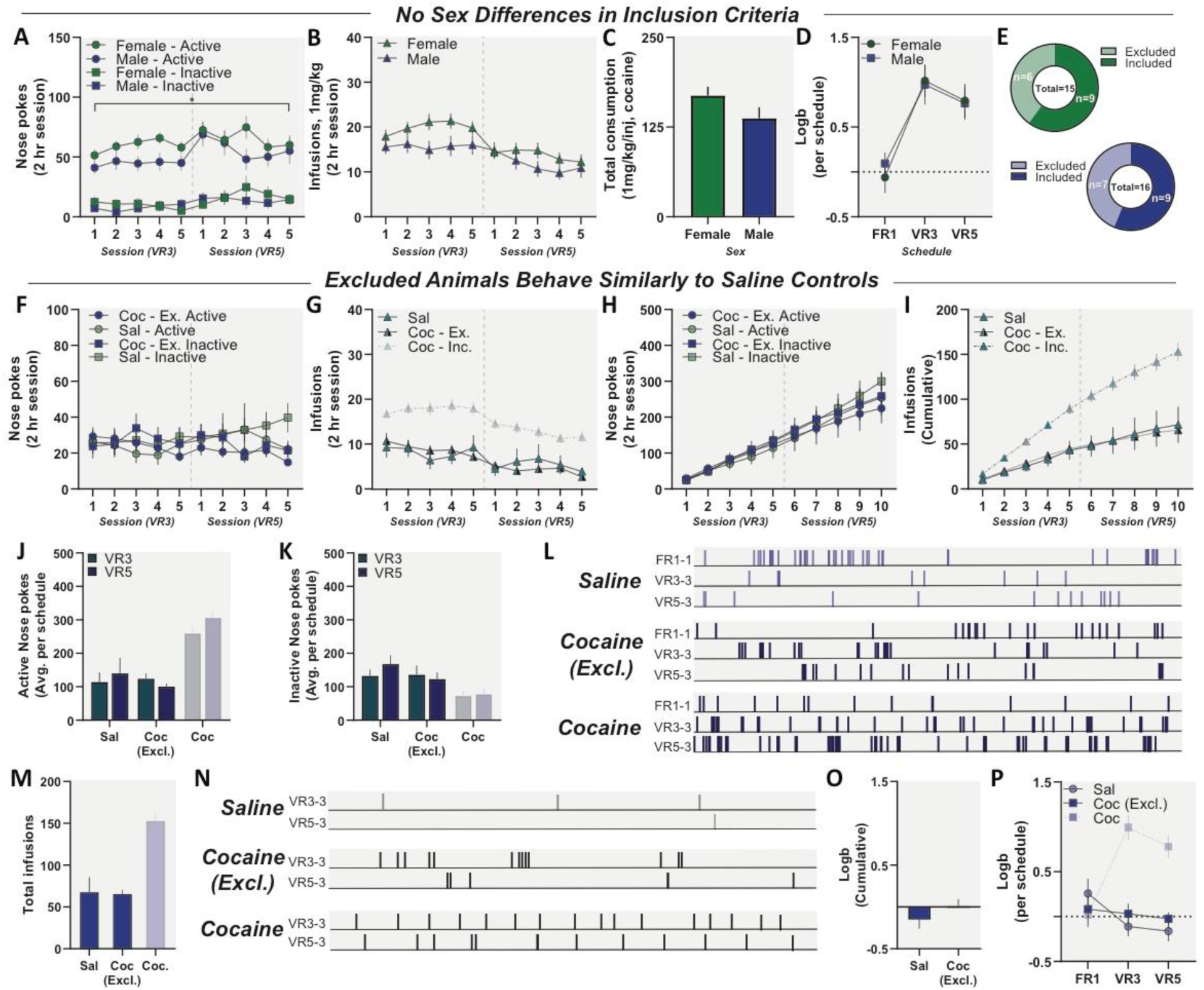
Variable-ratio-based inclusion criteria eliminates sex bias. [*Grey-scale: Cocaine-Included animals, excluded from subsequent analysis*](**A**) Active and inactive responses for female and male mice during cocaine self-administration. Male and female mice show similar preference for active nose-poke compared to inactive. (**B**) Cocaine infusions for female and male mice during cocaine self-administration. Female and male mice self-administer similar levels of cocaine. (**C**) Total cocaine consumption in self-administration (VR3 and VR5). Female and male mice consume similar levels of cocaine. (**D**) Female and male mice acquire bias for active nose-poke at similar rates during cocaine self-administration. (**E**) Female and male animals excluded based on VR criteria. VR criteria shows no sex bias in exclusion. (**F**) Active and inactive responses for Cocaine-Excluded and Saline animals during self-administration. Cocaine-Excluded animals show comparable active and inactive responses compared to Saline controls. (**G**) Infusions of cocaine or saline during self-administration. Cocaine-Excluded animals do not take more infusions than saline controls. (**H**) Cumulative active and inactive responses of Cocaine-Excluded animals during self-administration. (**I**) Cumulative infusions of Cocaine-Excluded and Saline animals during self-administration Cocaine-Excluded animals earn similar infusions to Saline controls. (**J**) Total active responses for Cocaine-Excluded animals and saline controls during VR3 and VR5. Cocaine-Excluded animals show no difference in active responding compared to saline controls. (**K**) Total inactive responses for Cocaine-Excluded animals and saline controls during VR3 and VR5. Cocaine-Excluded animals show no difference in inactive responding compared to saline controls. (**L**) Active response raster plots for representative (top) saline control, (middle) Cocaine-Excluded, and (bottom) Cocaine-Included animals. (**M**) Total infusions during cocaine self-administration, with no difference between Saline controls and Cocaine-Excluded animals. (**N**) Infusion raster plots for representative (top) saline control, (middle) Cocaine-Excluded, and (bottom) Cocaine-Included animals. (**O**) Saline and Cocaine-Excluded animals show no difference in response bias during self-administration. (**P**) Saline and Cocaine-Excluded animals do not acquire response bias for active nose-pokes throughout self-administration. Data reported as mean ± S.E.M. * p < 0.05

#### Training procedure

Animals were first trained on a FR1 schedule of reinforcement until ≧6 mg/kg of cocaine was consumed in a single session. These criteria were purely intake based and thus independent of responses on active pokes relative to inactive. Mice were subsequently trained on a series of VR schedules of reinforcement. VR schedules result in reinforcement following an unpredictable number of responses around an average number of responses per reinforcer delivery. VR schedules create steady and high rate of responding that exceed those of other fixed ratio or interval schedules[33–35]. Mice were switched to VR2 for 5 consecutive days or until criteria were met. Criteria were defined as 2 consecutive days of >70%, or 3 consecutive days of >60% discrimination (defined as the percentage of active versus total lever presses). Animals that fail to meet acquisition in 5 days were moved to VR3 self-administration and animals that fail to increase VR3 responding by >30% are excluded from studies. Following acquisition, animals were moved to self-administer under a VR3 schedule for 5 days and then to VR5 for an additional 5 days equaling 10 days of self-administration total. During VR3/5 self-administration, animals were excluded if they did not maintain >50% baseline infusions for 70% of self-administration sessions.

#### Analysis Parameters

For within-subject comparisons, a repeated measure ANOVA was used. For comparisons between groups, a two-tailed t-test was used. Each statistical test is denoted with the appropriate statistics in the results section. We also used a computational analysis to determine the parameters of response bias (Log *b*), as described previously[36–38]. Briefly, Log *b* was computed as the measure for behavioral bias using a logarithmic scale for the multiplication of the ratio between correct and incorrect responses: **Log *b = 0***.***5* * *log [(Active* + *0***.***5)***^***2***^***/(Inactive*+*0***.***5)***^***2***^]. Type I error rate (alpha) was set to 0.05 for all statistical tests. Data are represented as the mean +/- S.E.M in figures. Data were analyzed and graphed using Graphpad Prism 8.2 (La Jolla, CA).

## Results

### Current approaches for cocaine self-administration do not meet the criteria for volitional consumption

Drug self-administration training paradigms frequently begin with food pre-training to enhance acquisition and reduce attrition rates[7,11,17,25]. To determine if current approaches for self-administration were adequate, mice were first trained briefly to self-administer food with a single cue pairing under the control of a single active lever (**Fig. 1A**). Mice show higher responding on the active lever compared to inactive when food was delivered on an FR1 schedule, and active responding increased further when response requirements were raised to an FR2 schedule. (**Fig. 1B**; *main effect of Lever: F*_*(1,20)*_*=67*.*17, p*<*0*.*0001; main effect of Session: F*_*(5,99)*_*=11*.*83, p*<*0*.*0001; Interaction: F*_*(5,99)*_*=6*.*532, p*<*0*.*0001;* **Fig. 1D**). Mice show stable intake of sucrose pellets across sessions and schedules of reinforcement (**Fig. 1C**). Lastly, mice exhibited discrimination indices above threshold criteria typically used to determine acquisition (**Fig. 1E**; *One sample t-test [Theoretical mean = 70], FR1*_*1*_: *t*_*(10)*_*=0*.*04012, p=0*.*9688; FR1*_*2*_: *t*_*(10)*_*=3*.*133, p=0*.*0106; FR1*_*3*_: *t*_*(9)*_*=5*.*788, p=0*.*0003; FR2*_*1*_: *t*_*(10)*_*=3*.*994, p=0*.*0025; FR2*_*2*_: *t*_*(10)*_*=4*.*797, p=0*.*0007; FR2*_*3*_: *t*_*(10)*_*=3*.*319, p=0*.*0078*).

Next, we aimed to test if responding for the previously food-paired cue - i.e. conditioned reinforcement - was sufficient to maintain responding. Here, active responses resulted in only the presentation of the previously paired cue. Animals maintain high levels of active responding for the conditioned reinforcer alone (**Fig. 1F**; *main effect of Lever: F*_*(1,20)*_*=82*.*6, p*<*0*.*0001*). Mice also show no difference in cue deliveries over sessions (**Fig. 1G)** and maintained suprathreshold discrimination indices (**Fig. 1H**; *One-sample t-test [Hypothetical mean = 70], FR1*_*1*_: *t*_*(10)*_*=3*.*133, p=0*.*0106; FR1*_*2*_: *t*_*(9)*_*=5*.*788, p=0*.*0003; FR1*_*3*_: *t*_*(10)*_*=3*.*994, p=0*.*0025; FR1*_*4*_: *t*_*(10)*_*=4*.*797, p=0*.*0007; FR1*_*5*_: *t*_*(10)*_*=3*.*319, p=0*.*0078*). Active responding for the cue alone was not different from active responding for sucrose during the FR1 sessions (**Fig. 1I, Fig. 1J**). Thus, animals that undergo food pre-training continue to meet inclusion criteria for drug self-administration, even in the absence of positive reinforcers (such as food or drug). These results indicate that the standard food pretraining and acquisition criteria that are used in most drug self-administration studies in mice will not differentiate animals that are responding for conditioned reinforcement and those that are responding for drug.

### Conventional exclusion criteria generate a sex bias in behavioral selection

Conventional training parameters use a strict discrimination index as criteria for inclusion (**Fig. 2A, *left***). We applied such inclusion criteria to our previous food pre-trained animals and found a sex bias, where discrimination index criteria excludes more female than male mice (**Fig. 2A, *right***, *Two-tailed chi squared*, ***χ****2*_*(1)*_*=5*.*091, p=0*.*0241*). Although female and male mice maintain active responding across changing schedules of food reinforcement, female mice show increased inactive responding under FR2 (**Fig. 2B**; **Active lever:** *main effect of Session: F*_*(2*.*189, 25*.*83)*_*=7*.*016, p=0*.*0030;* **Inactive lever:** *main effect of Session: F*_*(2*.*215, 26*.*13)*_*=9*.*639, p=0*.*0005; main effect of Sex: F*_*(1,12)*_*=10*.*76, p=0*.*0066; Interaction: F*_*(5*.*59)*_*=9*.*115, p*<*0*.*0001*). Female-Excluded animals show a schedule-dependent decrease in discrimination index (**Fig. 2C**; *main effect of Session: F*_*(3*.*087,40*.*13)*_*=5*.*755, p=0*.*0021; main effect of Group: F*_*(2,65)*_*=39*.*23, p*<*0*.*0001; Interaction: F*_*(10,65)*_*=9*.*984, p*<*0*.*0001)*. Similarly, Sidak’s post-hoc analysis shows that Female-Excluded animals show lower discrimination as compared to Female-Included (t_(20.07)_=3.246, p=0.0121) and Male animals (t_(20.56)_=4.11, p=0.0016). However, despite sex-specific differences in inactive responding and discrimination index, all three groups maintain total reinforcers earned during food pre-training (**Fig. 2D**). Moreover, included animals maintain bias for active responding across training and test schedules, while Excluded-Females show a decrease in response bias (**Fig. 2E**; *main effect of Group: F*_*(2,11)*_*=15*.*03, p =0*.*0007; Interaction: F*_*(4,22)*_*=4*.*944, p=0*.*0054*). These data show that the higher rate of exclusion for female subjects using a traditional discrimination index is driven by increased responding on the inactive compared to males.

The traditional definition of reinforcement is a stimulus that maintains a rate of behavior that is higher than the unreinforced condition[35,39]. Thus, the more appropriate comparison in this case is baseline rates of responding in the absence of a reinforcer (**Fig. 2F, *left***). Indeed, Male, Female-Included, and Female-Excluded animals show increased responding above baseline under FR1 and FR2 schedules of reinforcement (**Fig. 2F, *right*;** *One-sample t-test [Hypothetical mean = 100]* **FR1:** *Male: t*_*(2)*_*=4*.*836, p=0*.*0402; Female-Included: t*_*(2)*_*=5*.*514, p=0*.*0314; Female-Excluded: t*_*(2)*_*=12*.*27, p=0*.*0066;* **FR2:** *Male: t*_*(2)*_*=5*.*639, p=0*.*0300; Female-Included: t*_*(2)*_*=12*.*81, p=0*.*0314; Female-Excluded: t*_*(2)*_*=7*.*444, p=0*.*0176)*. Lastly, there is no difference in responding when calculated as a percent of baseline (**Fig. 2G**). These data indicate that although Excluded-Females show high inactive responding, they meet the canonical definition of reinforcement and that strict discrimination indices select for male-typical behavior.

Similar to food pre-training, female and male animals maintain active responding for the cue alone under FR1 schedule of reinforcement as highlighted by a lack of statistical difference in cue responses over sessions (**Fig. 2H**). However, Female-Excluded animals demonstrate a decreased discrimination index during conditioned reinforcement testing (**Fig. 2I**; *main effect of Group: F*_*(2,11)*_*=15*.*91, p=0*.*0006)*. Sidak’s post-hoc test showed that Female-Excluded animals have lower discrimination indices compared to Female-Included mice (t_(19.97)_=5.469, p<0.0001) and males (t_(19.96)_=6.263, p<0.0001), while there was no difference between Female-Included and Male mice (t_(39.27)_=1.253, p=0.5213). However, despite decreases in discrimination index, Female-Excluded animals maintain stable levels of cue delivery where there are no differences in cues delivered between groups or over sessions (**Fig. 2J**). Thus, under traditional operant training and inclusion criteria, sex-biases are generated due to increased inactive responses by a subset of female mice, not due to decreases in acquisition rates.

### Variable-ratio reinforcement schedules engender high rates of responding and cocaine-intake

In previous work, response biases are generated by the inclusion of consequent stimuli (i.e. a light cue) only on the active operanda. Indeed, we have shown that use of cues in this manner paired with food pre-training in operant tasks can produce high discrimination indices in the absence of a primary reinforcer (see **Fig. 1F-J**). To circumvent these drawbacks, we used an operant schedule where active and inactive responses both generate programmed consequences (**Fig. 3A, *left***). Active responses result in delivery of both a positive reinforcer (i.e. a single cocaine infusion) and an active nose-poke-specific cue light, whereas inactive responses resulted only in the presentation of an inactive nose-poke-specific cue light (**Fig. 3A, *top right***). Further, we developed a novel operant training procedure that does not include food pre-training. Mice were trained to self-administer cocaine (1mg/kg/inj) under FR1 and VR2 reinforcement schedules until an adapted response criterion was met (**Fig. 3A, *bottom right***; see Methods for criteria). Mice then self-administered cocaine under VR3 for five consecutive days and under VR5 for five consecutive days. Following training, mice show a preference for active nose-poke compared to inactive that is schedule dependent (**Fig. 3B**; *main effect of Lever: F*_*(1,34)*_*=80*.*05, p*<*0*.*001; main effect of Session: F*_*(9,306)*_*=5*.*062, p*<*0*.*0001; see* **Fig. 3F**). Mice also maintain steady levels of cocaine intake, although show a slight decrease in intake under VR5 (**Fig. 3C**; *F*_*(4*.*763,80*.*97)*_*=11*.*81, p*<*0*.*0001; see* ***Fig. 3G***). Mice in the saline group show no preference for the active nose-poke and do not increase active responding from VR3 to VR5 (**Fig. 3D, Fig. 3H**). Saline infusions do not change over schedules of reinforcement, however, saline animals receive fewer infusions than cocaine (**Fig. 3E**; *main effect of Cocaine: F*_*(1,24)*_*=19*.*18, p=0*.*0002*). Cocaine animals demonstrate increased response rates on the active lever throughout cocaine self-administration compared to saline controls (**Fig. 3J**; *main effect of Schedule: F*_*(1*.*522,35*.*77)*_*=22*.*61, p*<*0*.*0001; main effect of Cocaine: F*_*(1,24)*_*=9*.*619, p=0*.*0049; Interaction: F*_*(2,47)*_*=16*.*75, p*<*0*.*0001*). Mice self-administering cocaine also demonstrate higher active responding and show an increase in active responding under VR5, while generating less inactive responses compared to saline controls (**Fig. 3K**; **Active Responding:** *main effect of Cocaine: F*_*(1,24)*_*=15*.*51, p=0*.*0006; main effect of Schedule: F*_*(1,24)*_*=6*.*416, p=0*.*0183;* **Fig. 3L**; **Inactive Responding** *main effect of Cocaine: F*_*(1,24)*_*=9*.*643, p=0*.*0048*). Lastly, mice self-administering cocaine demonstrate increased bias for active nose-poke during VR self-administration compared to saline controls that is acquired during self-administration training (**Fig. 3M**; *Two-tailed t-test, t*_*(24)*_*=4*.*897, p*<*0*.*0001;* **Fig. 3N**; *two-way ANOVA, main effect of Cocaine: F*_*(1,24)*_*=15*.*47, p=0*.*0006; Interaction: F*_*(2,48)*_*=18*.*41, p*<*0*.*0001*). Our novel dual-cue VR training paradigm allows for rapid acquisition of cocaine self-administration. Critically, saline animals do not demonstrate active response bias in the absence of a positive reinforcer, as seen with other operant training paradigms (see **Fig. 1**).

### Adapted VR schedule does not show sex-specific exclusion bias

Commonly used operant training paradigms generate sex-biases, as outlined in **Fig 2**. We assessed any sex-specific effects in our dual-cue VR training paradigm. During cocaine self-administration, there was no difference in active responses between female and male mice and no difference in inactive responses between female and male mice (**Fig. 4A**). Female and male mice received similar levels of cocaine infusions and comparable levels of total cocaine intake (**Fig. 4B, C**). In addition, female and male mice acquire bias for active responding at equal rates (**Fig. 4D**; *main effect of Schedule: F*_*(1*.*306,20*.*90)*_*=24*.*71, p*<*0*.*0001; no effect of Sex: F*_*(1,16)*_*=0*.*02026, p=0*.*8886*). Lastly, we found no sex-specific effects on our adapted exclusion criteria, as female and male mice are equally excluded (**Fig. 4E**).

In evaluating our VR inclusion criteria, it is critical to assess the behavioral profile of not only Cocaine-Included animals (**Fig. 3**), but also animals that fail to meet inclusion criteria (Cocaine-Excluded). Here, we find that the behavioral profile of Cocaine-Excluded animals is more comparable to saline controls. Cocaine-Excluded mice show no difference in active or inactive responding (**Fig. 4F, H**). Cocaine-Excluded animals do not take more infusions than saline controls (**Fig. 4G, I**). Cocaine-Excluded animals show no difference in total active or inactive responding throughout self-administration (**Fig. 4J, L, K**). With regard to reinforcers, Cocaine-Excluded animals show no increase in total infusions compared to saline controls (**Fig. 4M, N**). Lastly, Cocaine-Excluded animals do not show response bias throughout self-administration and do not acquire response bias over self-administration sessions (**Fig. 4O, P**).

These data demonstrate that our adapted VR inclusion criteria does not generate a sex bias in included animals. Moreover, animals that fail to meet inclusion criteria (Cocaine-Excluded) perform similarly to saline-controls.

## Discussion

Together, we show that traditional mouse models of self-administration result in high levels of operant responding in the absence of a reinforcer - with equivalent response rates between reinforced and non-reinforced sessions - demonstrating that it is impossible to determine when intake is volitional, or if animals are responding for the food-conditioned reinforcer and have not associated this action with the delivery of cocaine. Given that the core assumption of self-administration models is that the intake is voluntary, this poses a particularly large problem for studies outlining the mechanisms of contingent drug consumption. Here, we present a novel training procedure for operant drug self-administration that allows for rapid acquisition of drug self-administration without the need for food pre-training (**Fig. 3**). The addition of an inactive lever to include a programed consequence (e.g. schedule-dependent delivery of a cue light) prevents the generation of a response bias in the absence of a reinforcer. Further, our response criteria accounts for sex-specific differences in performance and eliminates sex-bias for animals excluded from further study (**Fig. 4E**). This procedure is simple to implement in mice, eliminates the confounds of previous training, and provides a number of independent measures that can be combined with novel circuit- and molecular-dissecting approaches. Moving forward, this will be a powerful tool to rigorously assess various drug-induced adaptations in molecular and circuit function.

Previous work has demonstrated that mice perform operant tasks for visual stimuli alone and will do so in a manner that meets various operant inclusion criteria, including active lever discrimination, schedule dependent changes in responding, and resistance to extinction[29,30,40]. As such, operant tasks programmed to deliver cues only to active responses can generate active-lever bias independent of reinforcer delivery. This is particularly problematic in drug self-administration, as active lever bias is often used as the read-out for not only volitional intake, but also catheter patency. In animals trained in single-cue paradigms, this active bias can be a product of the cue delivery and not drug-seeking. Behaviorally, this is relatively straight-forward to resolve, as conditioned reinforcement can be parsed from the acquisition of drug-self-administration via extinction of sucrose responding prior to training for intravenous drug infusions[6]. However, in experiments whose end goal is to assess the molecular and circuit adaptations induced by drug-seeking and drug-taking, a history of extinction training introduces a confound for subsequent drug-induced adaptations and altered circuit function [41,42]. As such, it is critical that operant procedures in mice take are designed to accurately induce and measure volitional drug intake independent of non-drug training.

A major component of addiction research has been characterizing the behavioral strategies and adaptations in drug-seeking and drug-taking paradigms[20,43–48]. A plurality of studies, however, focus only on the male phenotype, as female subjects are either underpowered or excluded altogether[49]. As a result, there has been a recent emphasis in accurately and reliably assessing sex as a biological variable with regard to preclinical models of substance use disorder[50–55]. Here, we assessed the sex-specific effects of food pre-training and commonly employed performance criteria for inclusion in subsequent operant studies. We show that typically used exclusion criteria, based on a discrimination index, generates a sex bias (**Fig. 2**). Specifically, female mice are more likely to be excluded based on decreased discrimination index compared to males (as a result of increased inactive responding), despite maintaining comparable rates of reinforcement and active responding. Traditional inclusion/exclusion criteria in mice rely on a strict discrimination index, where reinforcement is determined as a ratio of active to inactive responses, with the assumption that if reinforcement occurs that a majority of responses will be allocated to the active lever[4,5]. Yet, inactive responses have no programmed penalty, causing the exclusion of animals from self-administration studies that may demonstrate key features of volitional intake but have higher non-specific responding. Moreover, reinforcement is canonically defined rates of behavior that are higher for the reinforced over the non-reinforced condition (in the case of a drug – rates of responding for drug as compared to vehicle) [35,56]. Thus, inactive responses without programmed consequences should not factor into these definitions. Further, previous work has shown sex-differences in baseline and drug-induced motor activity[57–60], two factors in mice that can further interact with the above parameters to alter the behavioral read-out of self-administration. This is problematic as it not only selects for male-type behavioral profiles but can also mask sex-differences in subsequent drug-associated behaviors. Several studies have demonstrated that female and male mice employ unique behavioral strategies in operant reinforcement tasks[61]. As such, if sex is to be accurately considered as a biological variable in behavioral, circuit, and molecular analyses, training paradigms and selection criteria should be able to parse volitional intake while taking into account baseline behavioral differences between male and female subjects.

Our results highlight several problems with currently employed operant training procedures in mice. In addition, the novel procedure presented here relies on VR schedules of reinforcement to minimize these confounds. VR schedules result in reinforcement following an unpredictable number of responses around an average number of responses per reinforcer delivery. VR schedules are particularly powerful as they create steady and high rates of responding that exceed those of other fixed ratio or interval schedules[34,35,62]. Here, we show VR schedules minimize inactive responses, increase intake, and generate high rates of active responses allowing for a clear delineation between animals that have acquired the task and those that have not. Further, this schedule does not require food pre-training, allowing for drug-reinforcement that is not influenced by other factors and will be critical in this study to dissociate drug-reinforcer from natural-reinforcer induced changes. Finally, this reinforcement schedule allows for a large number of independent measures that will be able to be used to correlate with the neural measures in molecular and circuit-based studies.

The goal of preclinical SUD work is to understand drug-taking behavior on a mechanistic level that would allow for the development of novel pharmacotherapeutic targets for treatment in clinical populations[63–65]. In recent years, the field has focused on identifying the various molecular and circuity adaptations induced by drugs of abuse and underlying drug-associated behaviors[66–75]. However, our ability to effectively characterize the components driving drug-seeking and addictive phenotypes depend entirely on having translational and rigorous mouse models of volitional drug consumption. Here we identify the problems with the current models and create a new optimized procedure that minimizes these issues. Together this approach will allow for more rigorous studies that result in a more complete and definitive understanding of the behavior at the core of these translational models.

## Acknowledgements

We would like to thank the NIDA drug supply program for providing drugs used within this study.

